# PLCγ2 Controls Neutrophil Sensitivity through Calcium Oscillation and Gates Chemoattractant Concentration Range for Chemotaxis

**DOI:** 10.1101/2025.04.07.647573

**Authors:** Xuehua Xu, Woo Sung Kim, Arthur Lee, Tian Jin

## Abstract

The connection between calcium oscillation and cell sensitivity is poorly understood. Calcium oscillation is triggered either spontaneously or upon receptor-ligand binding. The cytosolic [Ca^2+^] increase during calcium oscillation is initiated from Ca^2+^ release from the intracellular stores through the phospholipase C (PLC)-derived inositol 1,4,5-trisphosphate (IP_3_). Here, we show that neutrophils lacking PLCγ2 (*plcg2^kd^*) display impaired spontaneous calcium oscillation and chemoattractant-induced calcium response, decreased membrane targeting of CAPRI (a RasGAP), and subsequent increased activations of Ras and its effectors, such as PI_3_Kγ activation and actin polymerization. More importantly, *plcg2^kd^* neutrophils sense and respond to chemoattractant at a subsensitive chemoattraction. Taken together, our results demonstrate that PLCγ2 mediates spontaneous calcium oscillation, contributes to chemoattractant-triggered calcium response, controls neutrophil sensitivity through membrane targeting of CAPRI, and gates chemoattractant concentration range for neutrophil chemotaxis.

**Highlights:**

- Neutrophils lacking PLCγ2 (*plcg2^kd^*) shows impaired spontaneous or GPCR-mediated Ca^2+^ oscillation.
- *plcg2^kd^* neutrophils display attenuated membrane targeting of CAPRI, a key negative regulator of Ras signaling and basal activity of neutrophils.
- *plcg2^kd^* neutrophils display an increased sensitivity and activation of GPCR-mediated signaling pathways.
- *plcg2^kd^* neutrophils display chemoattractant concentration-dependent alteration of chemotaxis behavior.

## INTRODUCTION

Calcium oscillation is ubiquitous, triggered either spontaneously or upon receptor-ligand binding in cells. The cytosolic [Ca^2+^] increase in calcium oscillation results from a coordinated release of intracellular Ca^2+^ stores and increased Ca^2+^ influx across the plasma membrane. The intracellular release of Ca^2+^ most commonly results from the Ca^2+^ release through the phospholipase C (PLC)-derived second messenger, inositol 1,4,5-trisphosphate (IP_3_), whereas the entry of Ca^2+^ is through the activation of store-operated channels (SOCs) in the plasma membrane. Chemoattractant-induced calcium oscillation, often also called calcium response, has been documented in neutrophils two decades ago (Petty, 2001). Chemoattractant-induced calcium response is significantly impaired in the mouse neutrophils deficient of PLCβ2/3, demonstrating the essential roles of PLCβ2/3 in chemoattractant-induced calcium response (Li et al., 2000). The potential connection between calcium oscillation and cell migration has been proposed (Clark et al., 2010; Petty, 2001). Local calcium pulse-mediated lamellipodia retraction and adhesion along the front of migration cells have also been reported (Tsai and Meyer, 2012). The essential role of PLCβ2/β3 in regulating the cofilin phosphatase slingshot 2 and neutrophil polarization and chemotaxis has been revealed (Tang et al., 2011). However, the player that mediates spontaneous calcium oscillation is not identified. The connection and relationship between spontaneous calcium oscillation and chemoattractants-induced calcium response remain unknown. More importantly, the biological function of the spontaneous calcium oscillation and the link between calcium oscillation and cell sensitivity remain elusive.

Phospholipase C gamma (PLCγ) is a potent regulator of many signaling pathways that are essential in physiological and pathological responses of immune disorders and cancers (Koss et al., 2014). Gain-of-function mutants of PLCγ2 have been the focus of investigations due to the patients developing several severe autoimmune or immunodeficiency conditions (Ombrello et al., 2012). The consequence of PLCγ2 deficiency gained more attention recently (Caraux et al., 2006; Jing et al., 2021; Obst et al., 2021; Palavicini et al., 2022). Contradictory conclusions have been made on the potential connection of down-regulations of PLC isoform and oncogenes Ras (Behjati et al., 2014; Danielsen et al., 2011; Martins et al., 2014). Neutrophils highly express PLCγ2, in addition to PLCβ2/β3 (Suh et al., 2008). Phosphorylation-mediated activation of PLCγ2 and its essential role in integrin/Fc receptor-mediated neutrophil functions have been reported (Jakus et al., 2009). Interestingly, we found that PLCγ2 is actively recruited to the leading edge of migrating neutrophils (Xu et al., 2015). Chemoattractant stimulation triggers a robust plasma membrane (PM) targeting of PLCγ2, an unconventional way of PLCγ activation (Falasca et al., 1998; Nishida et al., 2003; Xu et al., 2015). Importantly, *plcg2* stably knocked-down human neutrophil-like (*plcg2^kd^*) cells display an altered PLC signaling, including DAG production and IP_3_-mediated calcium response (Xu et al., 2023a), demonstrating the involvement of PLCγ2 in chemoattractant-mediated PLC signaling in neutrophils. Specifically, *plcg2^kd^*cells display a reduced duration of calcium response in response to chemoattractant stimulation at a saturating concentration. Neutrophils highly express CAPRI (<u>C</u>alcium-<u>p</u>romoted <u>R</u>as inactivator), a GAP protein that possesses a calcium-binding C2-domain and targets to plasma membrane upon cytosolic [Ca2^+^] increase (Nalefski and Falke, 1996). Thus, CAPRI is considered one of those Ras GAP proteins decoding calcium signaling for Ras activation (Liu et al., 2005). Consistent with the above, this altered calcium response results in a decreased PM targeting of CAPRI and subsequent hyper Ras activation and impaired polarization and chemotaxis of *plcg2^kd^*neutrophils (Lockyer et al., 2001; Xu et al., 2023b). We recently found that CAPRI controls the sensitivity of human neutrophils and gates the concentration range of chemoattractant gradients for long-range chemotaxis (Xu et al., 2021). Here, we investigated the role of PLCγ2 in spontaneous calcium oscillation and chemoattractant-induced calcium response for PM targeting of CAPRI and its effect on Ras activation and its downstream effectors in neutrophils. In conclusion, our results demonstrate that PLCγ2 mediates spontaneous calcium oscillation, contributes to chemoattractant-triggered calcium response, controls the sensitivity of neutrophils through membrane targeting of CAPRI, and gates chemoattractant concentration ranges for neutrophil chemotaxis.

## MATERIALS AND METHODS

### Cell culture and differentiation

The culturing of control and *plcg2kd* HL60 cells was as previously reported (Xu et al., 2023b). Briefly, cells were maintained in RPMI 1640 culture medium [RPMI 1640 medium with 20% (v/v) fetal bovine serum and 25 mM HEPES (Quality Biological, Inc. Gaithersburg, MD)]. HL60 cells were differentiated in RPMI 1640 culture medium containing 1.3% DMSO for 5 days before the experiments. The cells were incubated at 37°C in a humidified 5% CO_2_ atmosphere.

### Plasmids and transfection of cells

The DNA vectors of turboGFP (tGFP)-human CAPRI, active Ras sensor (active <u>R</u>as <u>b</u>inding <u>d</u>omain of human Raf1 tagged with mRFP, RBD-RFP), PIP_3_ biosensor (PH-GFP), and PM markers (CAAX-mCherry) were from Addgene (Cambridge, MA). The F-actin sensor F-tractin-GFP was obtained from John Hammer (Yi et al., 2012). The transfection procedure was as previously described (Xu et al., 2016). Briefly, 2 × 10^6^ cells were centrifuged at 100 × *g* for 10 min and resuspended in a mixture of 80 µL nucleofection solution V and 20 μL supplement I at room temperature. Six micrograms of plasmid DNA encoding the cDNA of the desired proteins were used for a single transfection reaction using program T-019 on the Amaxa Nucleofector II (Lonza, MD).

### Calcium response

Cells were incubated with 100 ng/ml Fluo4 (Invitrogen, Carlsbad, CA) at 37°C for 30 min, washed with RPMI 1640 medium with 25 mM HEPES twice to remove the unstained Fluo-4, and then subjected to the experiments.

### Ras and Rap1 activation assay

The procedure was as previously reported (48). Briefly, cells were starved with RPMI medium containing 25 mM HEPES at 37 ℃ for 3 hours. Cells were then collected and resuspended at 2×10^7^ cells/ml and then stimulated with a final concentration of 1 μM fMLP. Aliquots of the cells were taken at the indicated time points after stimulation and mixed with IB. The mixtures were incubated on ice for 30 min and then centrifuged at 100,000 × *g* at 4 ℃ for 30 min. The supernatants were incubated with agarose beads conjugated with RBD (active Ras binding domain of human Raf1) (Cytoskeleton, Inc. Denver, CO) or RBD_RalGDS_ (Rap1-GTP binding domain of human RalGDS) (abcam, Cambridge, UK) at 4 ℃ for 2 hours. The agarose beads were washed three times with IB. The protein on the beads was eluted by mixing with 25 μl SLB. The supernatants and eluted proteins were subjected to western blot detection of the indicated proteins.

### Imaging and data processing

Cells were plated and allowed to adhere to the cover glass of a 4-well or a 1-well chamber (Nalge Nunc International, Naperville, IL) precoated with Fibronectin (Sigma Aldrich, Saint Louis, MO) for 10 min, and then covered with RPMI 1640 medium with 10% FBS and 25 mM HEPES. For confocal microscopy, cells were imaged using a Carl Zeiss Laser Scanning Microscope Zen 780 (Carl Zeiss, Thornwood, NY) with a Plan-Apochromat 60x/1.4 Oil DIC M27 objective. For the uniform-stimulation experiment of membrane translocation assays, the stimuli were directly delivered to the cells as previously described (Xu et al., 2016). The membrane translocation of the indicated protein was measured by the depletion of the interested protein in the cytoplasm. The data obtained were further analyzed with Microsoft Office Excel (Redmond, WA). For quantitative analysis of membrane translocation dynamics of the indicated molecules, the cytosolic depletion of the indicated molecule was measured. Regions of interest (ROIs) in the cytoplasm (avoiding the nucleus area as much as possible) were within the cells throughout the time period of the measurements. The periphery of the cells was marked by the membrane markers. For data analysis, to normalize the effect of photobleaching during data acquisition, the intensity of ROIs in the cytoplasm was first divided by the intensity of whole cells at each given time point. To normalize the effect of morphological change during the time period, the above resulting data were divided by the intensity of ROIs in the PM marker channel. Lastly, the resulting data were divided by that at time 0 s; consequently, the relative intensity of any cells at time 0 s became 1. The graph of mean ± SD is shown.

### TAXIScan chemotaxis assay and data analysis

The procedure was as previously reported (Wen et al., 2016). Briefly, differentiated cells were loaded onto fibronectin-coated 4-µm EZ-TAXIScan chambers. The chemoattractants at the indicated concentrations were added to the other side of the well across the terrace that the cells chemotax through. The cells migrated for 30 min at 37 °C. Images were taken at 30-s intervals. For chemotaxis parameter measurements, 20 cells in each group were analyzed with DIAS software (Wessels et al., 1998). The bar graphs of chemotaxis parameters in mean and SD were plotted with Microsoft Office Excel (Redmond, WA).

## RESULTS

### Impaired spontaneous calcium oscillation and chemoattractant-induced calcium response in *plcg2^kd^* cells

Calcium oscillation is ubiquitous, occurring spontaneously or triggered upon receptor-ligand binding in cells. To distinguish these two different initiating mechanisms of [Ca^2+^] increase in cells, we refer to the non-ligand-induced [Ca^2+^] increase as spontaneous calcium oscillation (or calcium oscillation) and the ligand-induced one as calcium response in the present study. We have previously monitored calcium responses in response to fMLP stimulation in control (CTL) HL60 cells or *plcg2* stably knocked-down (*plcg2^kd^*) HL60 cells (Xu et al., 2023b). Upon a saturating fMLP stimulation, there is no significant difference in the amplitude of calcium response between CTL and *plcg2^kd^*cells. The difference is that, comparing CTL cells, *plcg2^kd^* cells display significantly reduced duration of calcium response and secondary sporadic calcium response. To have a better understanding of PLCγ2’s function in calcium response or spontaneous calcium oscillation, we monitored fMLP-induced calcium response in both CTL and *plcg2^kd^* cells at three different concentrations (10 nM, 1 nM, and 0.1 nM). To quantify calcium response, it is important to visualize the application of stimuli, especially the subsensitive ones. Thus, we mixed fMLP with a fluorescent dye, Alexa 633 (red), and simultaneously monitored the application of a mixture of fMLP stimuli (red) to the cells and the intensity change of the calcium indicator, fluo4 (green), in these cells (**Figure 1**). A clear calcium response is triggered in both CTL and *plcg2^kd^* cells in response to either 10 nM (**Figure 1A** and Video S1) or 1 nM fMLP stimulation (**Figure 1C** and Video S2). Compared to that of CTL cells, the amplitude and duration of calcium response in *plcg2^kd^* cells were significantly reduced (**Figure 1B** and **1E**), indicating that PLCγ2 contributes to both the amplitude and the duration of calcium responses upon stimuli at a relatively low concentration. Upon 0.1 nM fMLP stimulation (a subsensitive concentration for CTL cells in the previous report) (Xu et al., 2021), neither CTL nor *plcg2^kd^* cells show clearly synchronized calcium response (**Figure 1E** and Video S3) (Xu et al., 2021). Importantly, CTL cells display sporadic calcium responses and spontaneous calcium oscillation as mentioned previously (**Figure 1F**) (Petty, 2001). In contrast to CTL cells, *plcg2^kd^* cells rarely show sporadic calcium response, indicating that PLCγ2 mediates spontaneous calcium oscillation in neutrophils. Taken together, the above results demonstrate that PLCγ2 not only constitutes the GPCR-mediated calcium response but also mediates the spontaneous calcium oscillation in the resting neutrophils.

**Figure 1.**
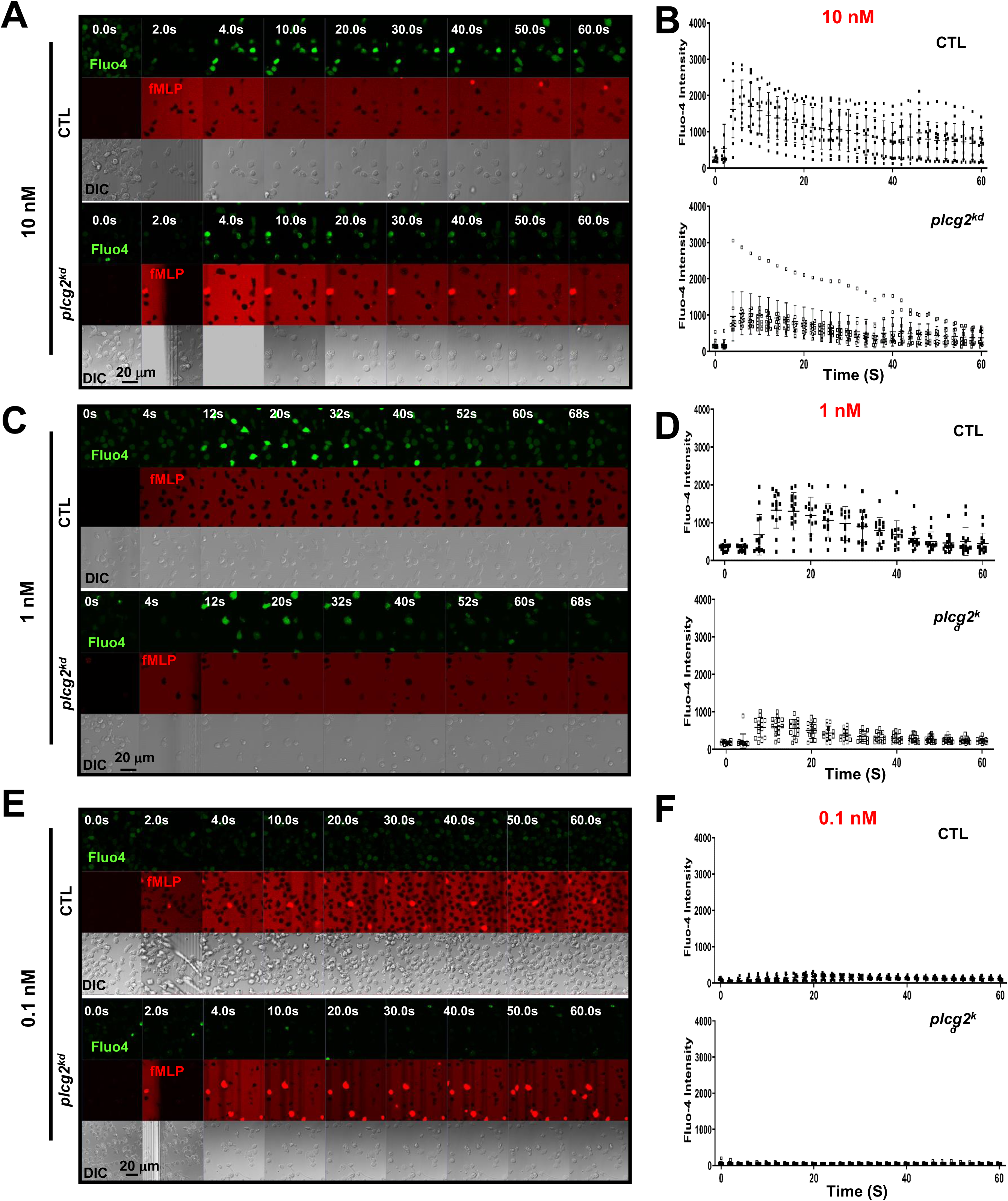
Decreased calcium response in *plcg2^kd^* cells upon fMLP stimulation. **A.** Montages show a 10 nM fMLP-induced calcium response in control (CTL) and *plcg2*-stably knocked down (*plcg2^kd^*) cells. Cells stained with the calcium indicator, Fluo-4 (green), were stimulated with fMLP at the indicated concentrations of fMLP at time 0 s. To visualize the application of fMLP stimuli, fMLP was mixed with a fluorescent dye, Alexa 633 (red). The parameters of image acquisition in CTL and *plcg2^kd^* cells are the same in **A**, **C**, and **D**. Scale bar = 20 μm. For complete sets of cell responses, see Video S1. CTL cells are in the upper panel, and *plcg2^kd^*cells are in the lower panel. The appearance of red indicates the application of fMLP stimulation. **B.** Dot plot analysis of Fluo-4 intensity change in CTL and *plcg2^kd^* cells before and after 10 nM fMLP stimulation in **A** and two other independent experiments. **C.** Montages show 1 nM fMLP-induced calcium response in CTL and *plcg2^kd^* cells. Cells stained with Fluo-4 (green) were stimulated with 1 nM fMLP at time 0 s. To visualize the application of fMLP stimuli, fMLP was mixed with a fluorescent dye, Alexa 633 (red). Scale bar = 20 μm. For complete sets of cell responses, see Video S2. CTL cells are in the upper panel, and *plcg2^kd^* cells are in the lower panel. The appearance of red indicates the application of fMLP stimulation. **D.** Dot plot analysis of Fluo-4 intensity change in CTL and *plcg2^kd^* cells before and after 1 nM fMLP stimulation in **C** and the other two independent experiments. **E.** Montages show 0.1 nM fMLP-induced calcium response in CTL and *plcg2^kd^* cells. For complete sets of cell responses, see Video S3. CTL cells are in the upper panel, and *plcg2^kd^* cells are in the lower panel. The appearance of red indicates the application of fMLP stimulation. **F.** Dot plot analysis of Fluo-4 intensity change in CTL and *plcg2^kd^* cells before and after 0.1 nM fMLP stimulation in **E** and two other independent experiments.

### Reduced membrane translocation of CAPRI in *plcg2^kd^* cells upon fMLP stimulations

The direct connection between calcium oscillation and cell sensitivity toward chemoattractant stimulation is not clear. We have previously shown that CAPRI mediates the deactivation of the GPCR-mediated Ras signaling to achieve Ras adaptation in human neutrophils (Xu et al., 2021). In resting neutrophils, the majority of CAPRI is cytosolic. However, a fraction of CAPRI localizes in the plasma membrane (PM) where it regulates basal Ras activity to control cell sensitivity. The PM targeting of CAPRI requires its C2-domain and proper [Ca^2+^] increase. However, it was not clear whether PLCγ2 is involved to mediate the [Ca^2+^] increase to regulate CAPRI membrane targeting in the resting cells. As previously reported (Xu et al., 2021), migrating CTL cells actively recruit CAPRI-GFP to the leading fronts, while *plcg2^kd^* cells rarely show a clear CAPRI-GFP localization on the protrusion sites (**Figure 2A**). Upon 10 nM fMLP stimulation, CTL cells display a robust membrane translocation of CAPRI-GFP (**Figure 2A**, upper panel, and Video S4, upper panel). However, *plcg2^kd^* cells display significantly reduced PM translocation of CAPRI-GFP (Video S4, lower panel), consistent with the previous report (Xu et al., 2023b). Upon 0.1 nM fMLP stimulation, both CTL and *plcg2^kd^* cells do not display visible membrane translocation of CAPRI-GFP (**Figure 2A** and Video S5). Migrating CTL cells recruit CAPRI-GFP in the leading fronts, while *plcg2^kd^* cells do not. Quantitative measurement of CAPRI PM translocation in multiple cells confirms the above observations (**Figure 2B** and **2D**). Taken together, the above result indicates that *plcg2^kd^* cells display impaired membrane targeting of CAPRI in both resting and stimulated states.

**Figure 2.**
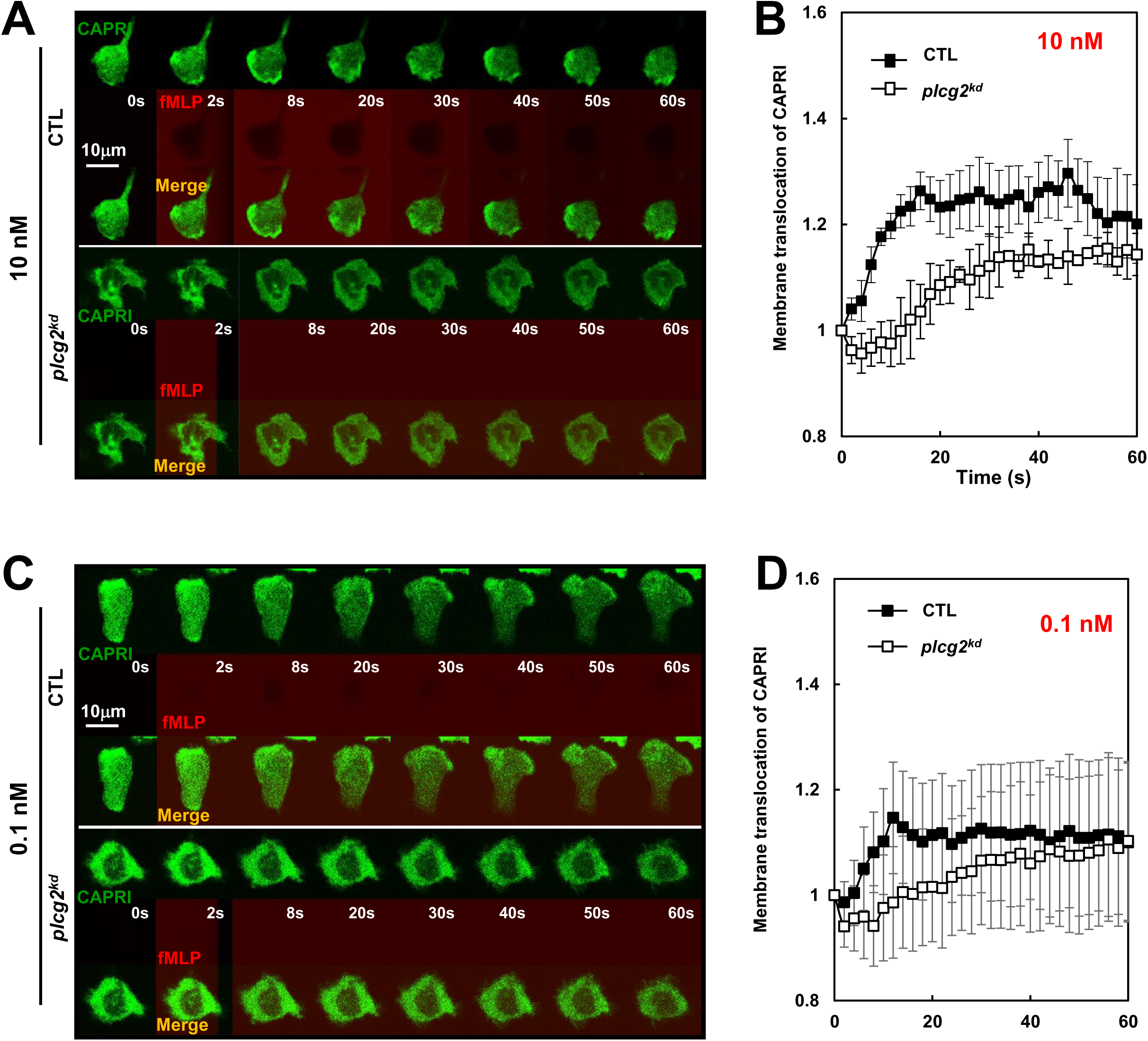
Reduced membrane translocation of CAPRI in *plcg2^kd^* cells upon fMLP stimulations. **A.** Montages show fMLP-induced plasma membrane (PM) translocation of CAPRI-GFP in CTL and *plcg2^kd^* cells in response to fMLP stimulation at 10 nM. Cells expressing CAPRI-GFP (green) were stimulated with fMLP stimulation at 10 nM at time 0 s. To visualize the application of the stimuli, fMLP was mixed with fluorescent dye, Alexa 633 (red). Scale bar = 20 μm. See Video S4 for complete sets of cell responses. The CTL cell is in the upper panel, and the *plcg2^kd^* cell is in the lower panel. **B.** Quantitative measurement of the membrane translocation of CAPRI-GFP in CTL and *plcg2^kd^* cells in response to 10 nM fMLP stimulation. Mean ± SD is shown; n = 5 or 5 for CTL or *plcg2^kd^* cells, respectively. **C.** Montages show fMLP-induced plasma membrane (PM) translocation of CAPRI-GFP in CTL and *plcg2^kd^*cells in response to fMLP stimulation at 0.1 nM. Cells expressing CAPRI-GFP (green) were stimulated with fMLP stimulation at 0.1 nM at time 0 s. To visualize the application of the stimuli, fMLP was mixed with fluorescent dye, Alexa 633 (red). Scale bar = 20 μm. See Video S5 for a complete set of cell responses. CTL cell is in the upper panel and *plcg2^kd^* cell is in the lower panels. **D.** Quantitative measurement of the membrane translocation of CAPRI-GFP in CTL and *plcg2^kd^* cells in response to 0.1 nM fMLP stimulation. Mean ± SD is shown; n = 5 or 5 for CTL or *plcg2^kd^* cells, respectively.

### Increased Ras activation in *plcg2^kd^* cells in response to fMLP stimulation at a low or a subsensitive concentration

To investigate the effect of impaired PM targeting of CAPRI in *plcg2^kd^*cells, we biochemically determined Ras activation using a pull-down assay in a large population of both CTL and *plcg2^kd^* cells upon fMLP stimulation at different concentrations as previously reported (Xu et al., 2021). 10 nM fMLP stimulation triggered a clear Ras activation in CTL cells and a stronger one in *plcg2^kd^*cells (**Figure 3A**-**3B**). 0.1 nM fMLP stimulation does not trigger a notable Ras activation in CTL cells, while it induces a clear Ras activation in *plcg2^kd^* cells, indicating the increased sensitivity in the cells lacking PLCγ2. To confirm the increased sensitivity of *plcg2^kd^* cells, we next monitored the temporospatial Ras activation using an active Ras probe (RBD-RFP, red) in both CTL and *plcg2^kd^* cells by confocal microscopy (**Figure 3C**). To visualize the application of the stimuli, fMLP was mixed with a fluorescent dye, Alexa488 (green). We found that RBD-RFP localizes at the protrusion site of both resting CTL and *plcg2^kd^* cells (Video S6). fMLP at 10 nM triggers a clear PM translocation of RBD-RFP in CTL cells and a significantly stronger and profoundly longer PM translocation of RBD-RFP in *plcg2^kd^*cells, consistent with the previous report (**Figure 3C**, upper panel, and Video S6) (Xu et al., 2023b). Upon 0.1 nM fMLP stimulation, the CTL cell does not display a clear overall PM translocation of RBD-RFP, instead of a continuous localization of RBD-RFP on the protrusion site or the leading front of a migrating cell (**Figure 3C**, lower panel, and Video S7, upper panel). However, the *plcg2^kd^* cell displays a clear PM translocation and subsequent continuous localization on the expanding protrusion sites of RBD-RFP upon the same 0.1 nM fMLP stimulation (Video S7, lower panel). Quantitative measurement of RBD-RFP membrane translocation in multiple cells further confirms the above observation (**Figure 3D**). In conclusion, *plcg2^kd^*cells display an increased Ras activation upon chemoattractant stimuli and responds to chemoattractant stimuli at a subsensitive concentration for CTL cells. The above result demonstrates that PLCγ2 not only is required for chemoattractant induced Ras adaptation but also controls cell sensitivity through PM-targeting of CAPRI in neutrophils.

**Figure 3.**
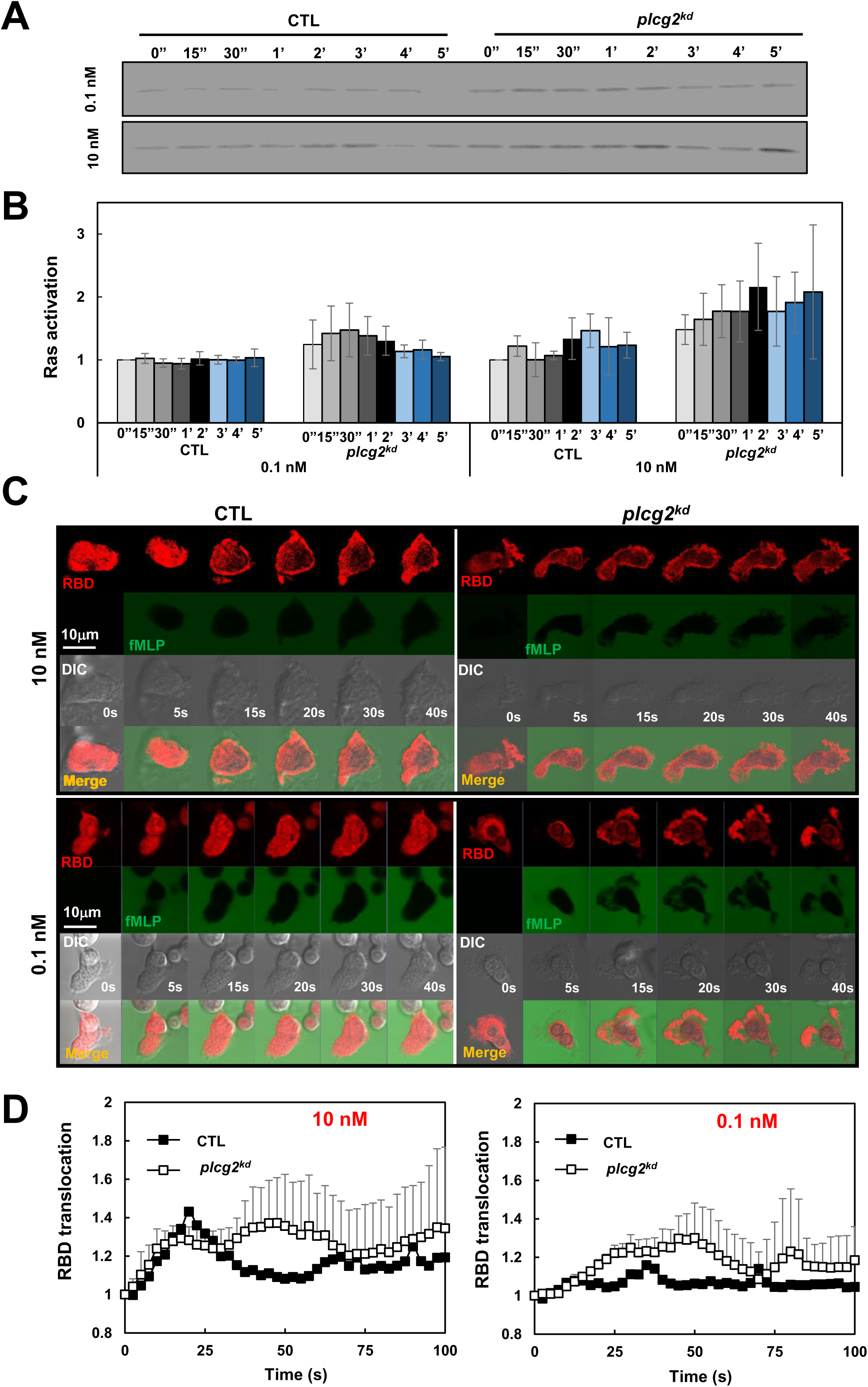
Increased Ras activation in *plcg2^kd^* cells in response to fMLP stimulation at a low (10 nM) or a subsensitive (0.1 nM) concentration of fMLP. **A.** Ras activation in CTL and *plcg2^kd^* cells in response to either 10 nM or 0.1 nM fMLP stimulation was determined by a pull-down assay. **B.** Normalized quantitative densitometry of the active Ras from three independent experiments, including the result presented in **A**. The intensity of active Ras in CTL cells at time 0 s was normalized to 1. Mean ± SD from the three independent experiments is shown. **C.** Montage shows fMLP-induced Ras activation in CTL (left) and *plcg2^kd^* (right) cells by the membrane translocation of the active Ras biosensor RBD-RFP. Cells expressing RBD-RFP (red) were stimulated with 10 nM (upper panel) or 0.1 nM (lower panel) fMLP at time 0 s. To visualize the application of fMLP, it was mixed with a fluorescent dye, Alexa 488 (green). Scale bar = 10 μm. See Video S6 or S7 (CTL, left panel; *plcg2^kd^*, right panel) for complete sets of cell responses upon fMLP stimulation at 10 nM or 0.1 nM, respectively. **D.** Quantitative measurement of PM translocation of RBD-RFP in CTL and *plcg2^kd^* cells in response to fMLP stimulation at either 10 nM (left) or 0.1 nM (right). Mean ± SD is shown. N = 4 or 4 for CTL or *plcg2^kd^* cells, respectively, in both graphs.

### Increased PI_3_K activation in *plcg2^kd^* cells in response to fMLP stimulation at a low or a subsensitive concentration

PI_3_Kγ, a direct effector of Ras, catalyzes the conversion of phosphatidylinositol (4,5)-bisphosphate (PI(4,5)P2, PIP_2_) to phosphatidylinositol (3,4,5)-trisphosphate (PtdIns(3,4,5)P3, PIP_3_) (Li et al., 2000; Suire et al., 2006; Suire et al., 2012). PI3Kγ activation recruits and activates the PIP_3_-binding serine/threonine kinase Akt on the plasma membrane, which plays an essential role in neutrophil chemotaxis (Tang et al., 2011). To examine the consequences of the increased Ras activation in *plcg2^kd^* cells, we next investigated PI_3_K activation in both CTL and *plcg2^kd^* cells. To determine PI_3_K activation, we monitored the production of PIP_3_ using a PIP_3_ biosensor, PH-GFP, in living HL60 cells by confocal microscopy. In the resting cells, PH-GFP (green) localizes mostly in the cytosol (**Figure 4A**). It also colocalizes with a plasma membrane (PM) marker (red) on the protrusion sites, more obvious in *plcg2^kd^*cells. Upon 10 nM fMLP stimulation, CTL cells display a transient translocation of PH-GFP to the periphery, where PH-GFP colocalizes with PM marker, followed by a continuous colocalization of PH-GFP and PM marker in the protruding sites (upper panel **in Figure 4A** and Video S8). Upon the same stimulation, the *plcg2^kd^* cell shows a significantly stronger and profoundly longer translocation of PH-GFP to the plasma membrane (lower panel in **Figure 4A** and Video S8). Quantitative measurement of RBD-RFP membrane translocation in multiple cells further confirms the above observation (**Figure 4B**). Upon 0.1 nM fMLP stimulation, the CTL cell does not display a clear overall PM translocation of PH-GFP, instead of a continuous localization on the protrusion site of a migrating cell (upper panel in **Figure 4C** and Video S9). However, upon the same 0.1 nM fMLP stimulation, *plcg2^kd^* cells display a clear PM translocation of RBD-RFP and subsequent continuous localization in the expanding protrusion sites (lower panel in **Figure 4C** and Video S9, and **Figure 4D**). In conclusion, *plcg2^kd^* cells display an elevated PI_3_K activation and can respond to chemoattractant stimuli at a subsensitive concentration.

**Figure 4.**
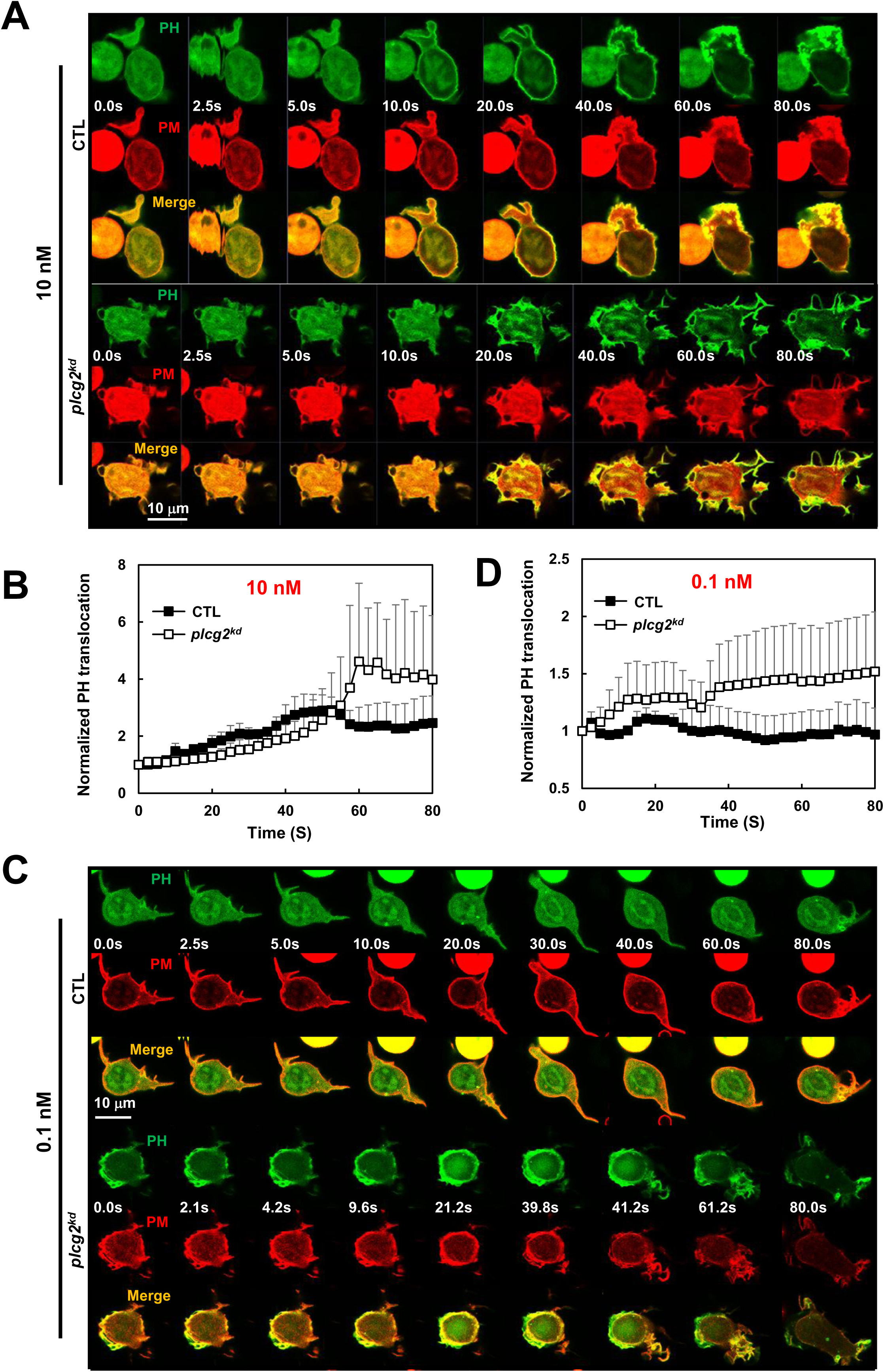
Increased PI_3_K activation in *plcg2^kd^* cells in response to fMLP stimulation at a low (10 nM) or a subsensitive (0.1 nM) concentration. **A.** Montage shows PI_3_K activation in CTL and *plcg2^kd^* cells in response to 10 nM fMLP stimulation by monitoring PIP_3_ production using fluorescent microscopy. PIP_3_ production is visualized by the membrane translocation of the PIP_3_ biosensor PH-GFP. Cells expressing PH-GFP (green) and a PM marker (red) were stimulated with 10 nM fMLP at time 0 s. Scale bar = 10 μm. See Video S8 (CTL, upper panel; *plcg2^kd^*, lower panel) for complete sets of cell responses upon 10 nM fMLP, respectively. **B.** Quantitative measurement of PIP_3_ production by the membrane translocation of PH-GFP in CTL and *plcg2^kd^* cells in response to 10 nM fMLP stimulation. Mean ± SD is shown; n = 3 or 5 for CTL or *plcg2^kd^* cells, respectively. **C.** Montage shows PI_3_K activation in CTL and *plcg2^kd^* cells in response to 0.1 nM fMLP stimulation by monitoring PIP_3_ production using fluorescent microscopy. Cells expressing PH-GFP (green) and a PM marker (red) were stimulated with 0.1 nM fMLP at time 0 s. Scale bar = 10 μm. See Video S9 (CTL, upper panel; *plcg2^kd^*, lower panel) for complete sets of cell responses upon 0.1 nM fMLP stimulation. **D.** Quantitative measurement of PIP_3_ production by the membrane translocation of PH-GFP in CTL and *plcg2^kd^* cells in response to 10 nM fMLP stimulation. Mean ± SD is shown; n = 5 or 5 for CTL or *plcg2^kd^* cells, respectively.

### Increased actin polymerization in *plcg2^kd^* cells in response to fMLP stimulation at a low or a subsensitive concentration

Neutrophils employ GPCR/G protein complexes to regulate multiple signaling pathways that control the actin cytoskeleton dynamics and drive cell migration. To evaluate the role of PLCγ2 in chemoattractant GPCR-mediated actin assembly in neutrophils, we monitored actin polymerization using the membrane translocation of an F-actin filament probe, F-tractin–GFP (green), in live cells by fluorescence microscopy (**Figure 5**). In resting cells, F-tractin-GFP (green) localizes in the cytosol and cortex, where it colocalizes with the PM marker (red) on the plasma membrane and protrusion sites (**Figure 5A**). We found that upon 10 nM fMLP stimulation at 2 s, more F-tractin-GFP translocated to the cell cortex at around 10 to 40 s, then mostly returned to the cytosol at about 60 s, and then translocated to the leading front again at around 80 s in CTL cells (upper panel in **Figure 5A** and Video S10). In response to the same 10 nM fMLP stimulation, *plcg2^kd^* cells displayed a continuous, persistent translocation of F-tractin-GFP and colocalized with the PM marker on the plasma membrane (lower panel in **Figure 5A** and Video S10). We further quantified the actin polarization of CTL and *plcg2^kd^*cells by the membrane translocation of F-tractin-GFP and confirmed that *plcg2^kd^*cells display elevated and prolonged actin polymerization compared to CTL cells (**Figure 5B**). We further determined the actin polymerization of both CTL and *plcg2^kd^*cells to 0.1 nM fMLP stimulation. In response to 0.1-nM fMLP stimulation, most CTL cells (∼90%) did not show the clear membrane translocation of F-tractin–GFP to the PM, while they showed cortex localization of F-tractin and PM marker (upper panel in **Figure 5C** and Video S11). In contrast, more than 80% of *plcg2^kd^* cells showed the clear membrane translocation of F-tractin-GFP upon 0.1 nM fMLP stimulation (lower panel in **Figure 5C** and Video S11). Quantitative measurement of membrane translocation of F-tractin in CTL and *plcg2^kd^* cells shows a normal oscillation of actin polymerization in CTL cells, while a clear actin polymerization in *plcg2^kd^* cells upon 0.1 nM fMLP stimulation (**Figure 5D**). These results together demonstrate that *plcg2^kd^* cells can mediate actin polymerization to chemoattractant stimulation at a subsensitive concentration, while they display a prolonged, elevated actin polymerization upon chemoattractant stimulation at low concentrations.

**Figure 5.**
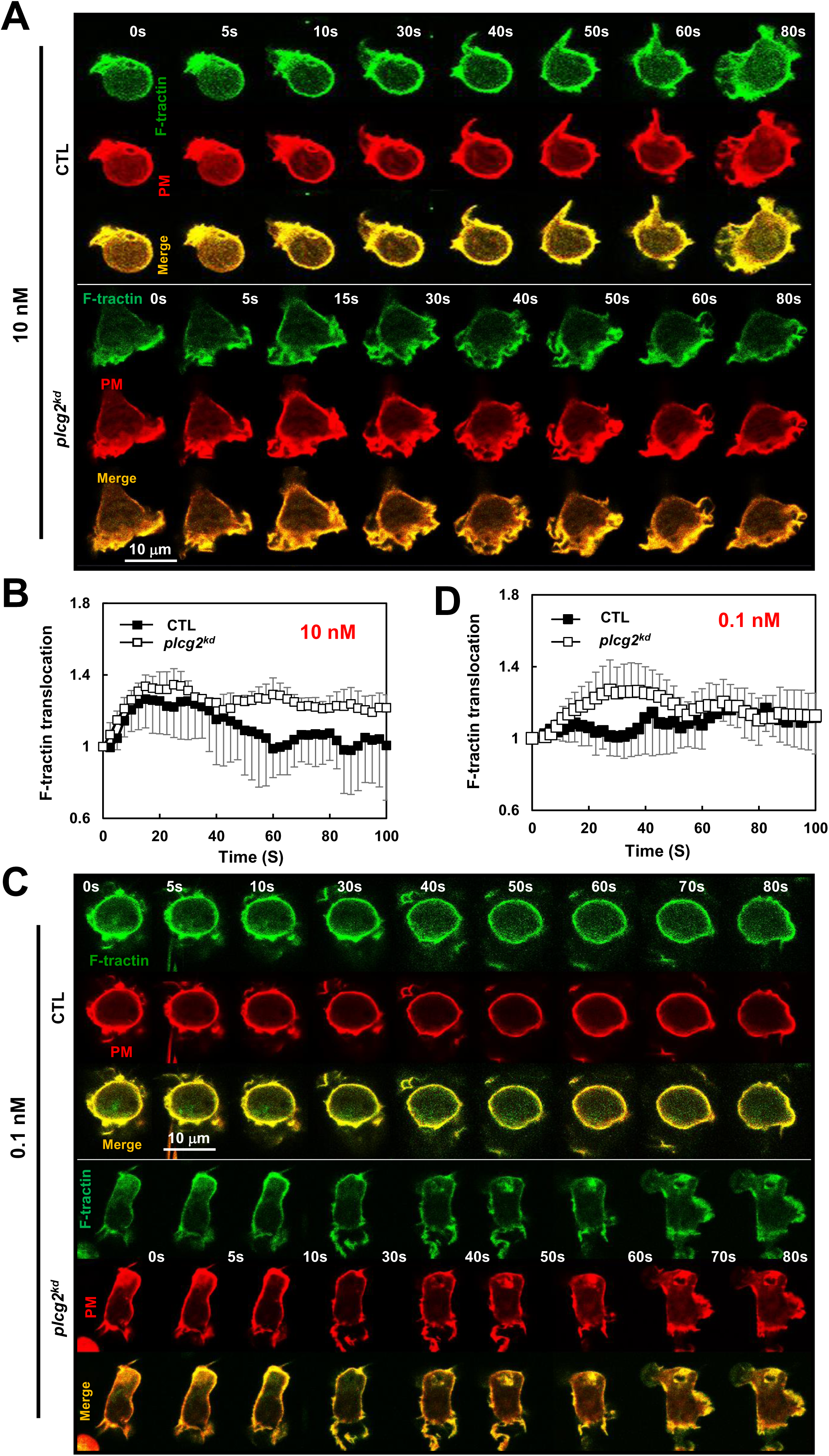
Increased actin polymerization in *plcg2^kd^* cells in response to fMLP stimulation at a low (10 nM) or a subsensitive (0.1 nM) concentration. **A.** Montage shows the membrane translocation of the F-actin probe (GFP-tagged F-tractin) in CTL and *plcg2^kd^*cells upon fMLP stimulation at 10 nM. Cells expressing F-tractin GFP (green) and a PM marker (red) were stimulated with fMLP at time 0 s. Scale bar = 10 μm. See Video S10 (CTL, upper panel; *plcg2^kd^*, lower panel) for a complete set of cell responses upon 10 nM fMLP stimulation. **B.** Quantitative measurement of actin polymerization by the membrane translocation of Ftractin-GFP in CTL and *plcg2^kd^* cells in response to fMLP stimulation. Mean ± SD is shown; n = 3 or 5 for CTL or *plcg2^kd^* cells, respectively. **C.** Montage shows the membrane translocation of the F-actin probe (GFP-tagged F-tractin) in CTL and *plcg2^kd^*cells upon fMLP stimulation at 0.1 nM. Cells expressing F-tractin GFP (green) and a PM marker (red) were stimulated with fMLP at time 0 s. Scale bar = 10 μm. See Video S11 (CTL, upper panel; *plcg2^kd^*, lower panel) for a complete set of cell responses upon 0.1 nM fMLP stimulation. **D.** Quantitative measurement of actin polymerization by the membrane translocation of Ftractin-GFP in CTL and *plcg2^kd^* cells in response to 0.1 nM fMLP stimulation. Mean ± SD is shown; n = 3 or 5 for CTL or *plcg2^kd^* cells, respectively.

### *plcg2^kd^* neutrophils chemotax in chemoattractant gradients at subsensitive concentrations

We found that *capri^kd^* neutrophils, which lack Ras inhibitor CAPRI, display an increased sensitivity and elevated activation of Ras and its downstream effectors (Xu et al., 2021). More importantly, *capri^kd^* neutrophils display an altered chemotaxis behavior: an improved chemotaxis in the gradients at subsensitive concentrations, a normal chemotaxis in the gradients at medium concentrations, and an impaired chemotaxis in the gradients at saturating concentrations. That is, neutrophils lacking CAPRI display an upshift in concentration range for chemotaxis (Xu and Jin, 2022). We have previously shown an impaired chemotaxis of *plcg2^kd^*cells upon chemoattractant gradients at a saturating concentration (Xu et al., 2023b). Next, we examined the chemotaxis behavior of CTL and *plcg2^kd^*cells in the gradients of three chemoattractants at either medium (100 nM) or subsensitive (0.1 nM) concentrations (**Figure 6** and Video S12). Without a gradient, *plcg2^kd^* cells displayed a better random walk compared to CTL cells (**Figure 6A-6B**), consistent with the previous report (Xu et al., 2023b). When exposed to gradients generated from a source at medium concentrations (100 nM), CTL and *plcg2^kd^* cells displayed overall similar chemotaxis capability, although *plcg2^kd^*cells displayed slightly decreased speed and total path length (**Figure 6B**). In the gradients generated from the sources of 0.1 nM, most CTL cells displayed random migration, while most *plcg2^kd^* cells displayed a clear directed cell migrating along the direction of the gradient. In conclusion, *plcg2^kd^*cells display a concentration-dependent, altered chemotaxis behavior: an improved chemotaxis in the gradients at a subsensitive concentration, while a normal chemotaxis in the gradients at medium concentrations. Combined with the previous report (Xu et al., 2023b), our results demonstrate that neutrophils lacking PLCγ2 display an upshift of the concentration ranges of diverse chemoattractants for an efficient chemotaxis.

**Figure 6.**
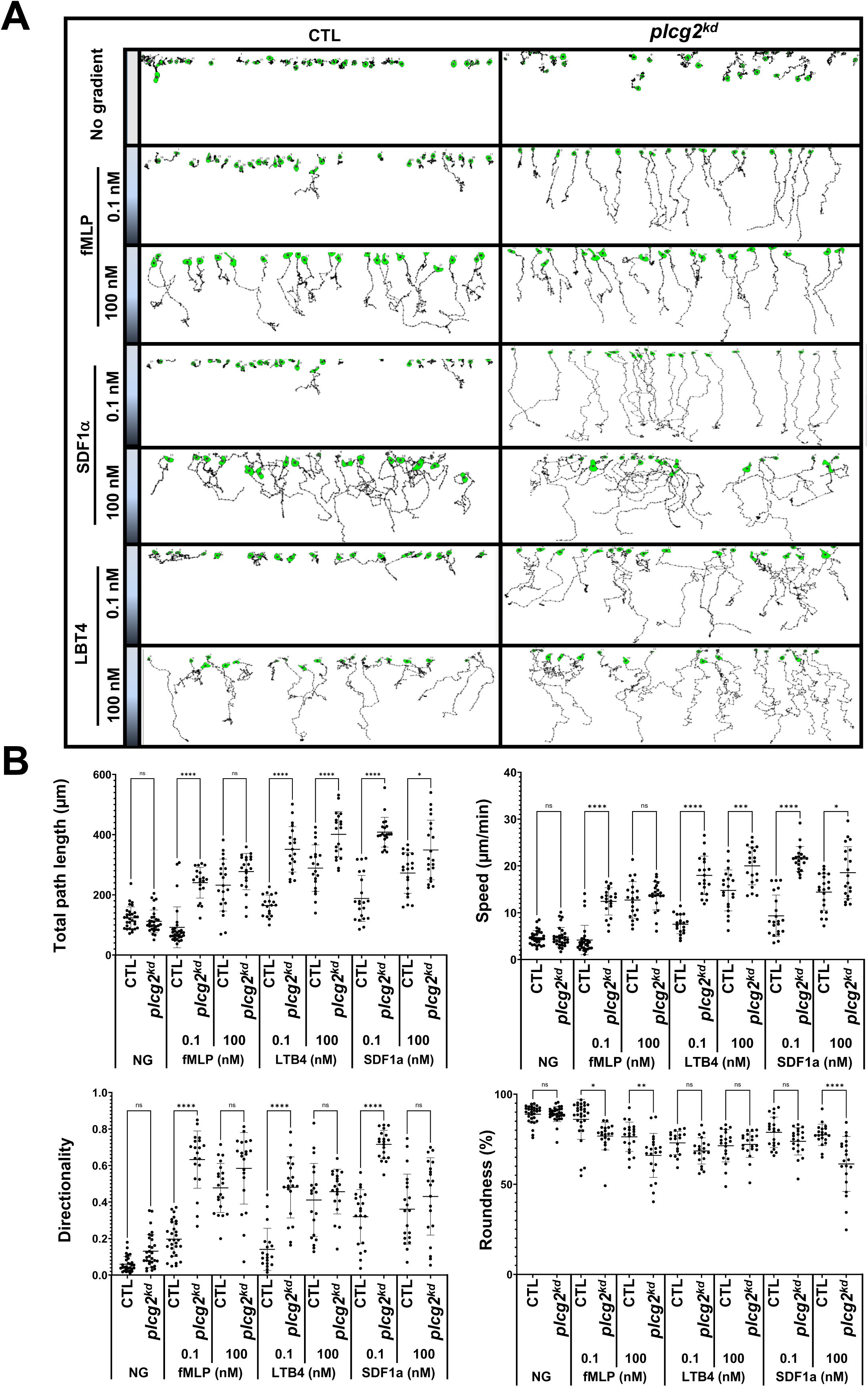
*plcg2^kd^* neutrophils display improved chemotaxis in chemoattractant gradients at subsensitive concentrations. **A.** Montages show the travel path of chemotaxing CTL or *plcg2^kd^* cells in response to subsensitive or mid-concentration gradients. The plain or shaded panels on the left side of the images in the montage indicate either no gradient (NG) or chemoattractant gradients of fMLP (top), SDF1a (middle), or LTB4 (bottom) sourced from the indicated concentrations. The concentration on the top side of the terrace is 0 and the concentration at the bottom side of the terrace is as indicated on the left side of the terrace. Movement of at least 30 cells in each group was analyzed by DIAS software and is shown. See Video S12 (CTL, left, and *plcg2^kd^*, right) for a complete set of ez-taxiscan images with the same conditions of chemoattractant concentrations shown in **A**. **B.** Chemotaxis behaviors measured from A are described as four parameters: directionality, which is “upward” directionality, where 0 represents random movement and 1 represents straight movement toward the gradient; speed, defined as the distance that the centroid of the cell moves as a function of time; total path length, the total distance the cell has traveled; and roundness (%) for polarization, which is calculated as the ratio of the width to the length of the cell. Thus, a circle (no polarization) is 1, and a line (perfect polarization) is 0. Thirty cells from each group were measured for 10 min. Student’s *t*-test was used to calculate the *p* values, which are indicated as *ns* (not significant *p* > 0.1), * (*p* < 0.1), ** (*p* < 0.01), *** (*p* < 0.001), or **** (*p* < 0.0001).

## DISCUSSION

Chemoattractant-triggered PLCβ2/β3 activation and the essential role of PLCβ2/β3 in subsequent calcium signaling in neutrophils have been previously characterized (Li et al., 2000). However, no connection has been made between calcium oscillation and cell sensitivity toward chemoattractants. The mediator(s) or biological functions of calcium oscillation in neutrophils largely remain elusive. In the present study, we show that PLCγ2-mediated spontaneous calcium oscillation controls the basal Ras activity and neutrophil sensitivity by the recruitment of CAPRI to the plasma membrane. More importantly, the chemoattractant-induced PLCγ2 activation constitutes the essential calcium response for the PM recruitment of CAPRI and subsequent adaptation of Ras and downstream effectors for proper chemotaxis. Hence, by applying the above two mechanisms, PLCγ2 gates the chemoattractant concentration range for neutrophil chemotaxis (**Figure 7**).

**Figure 7.**
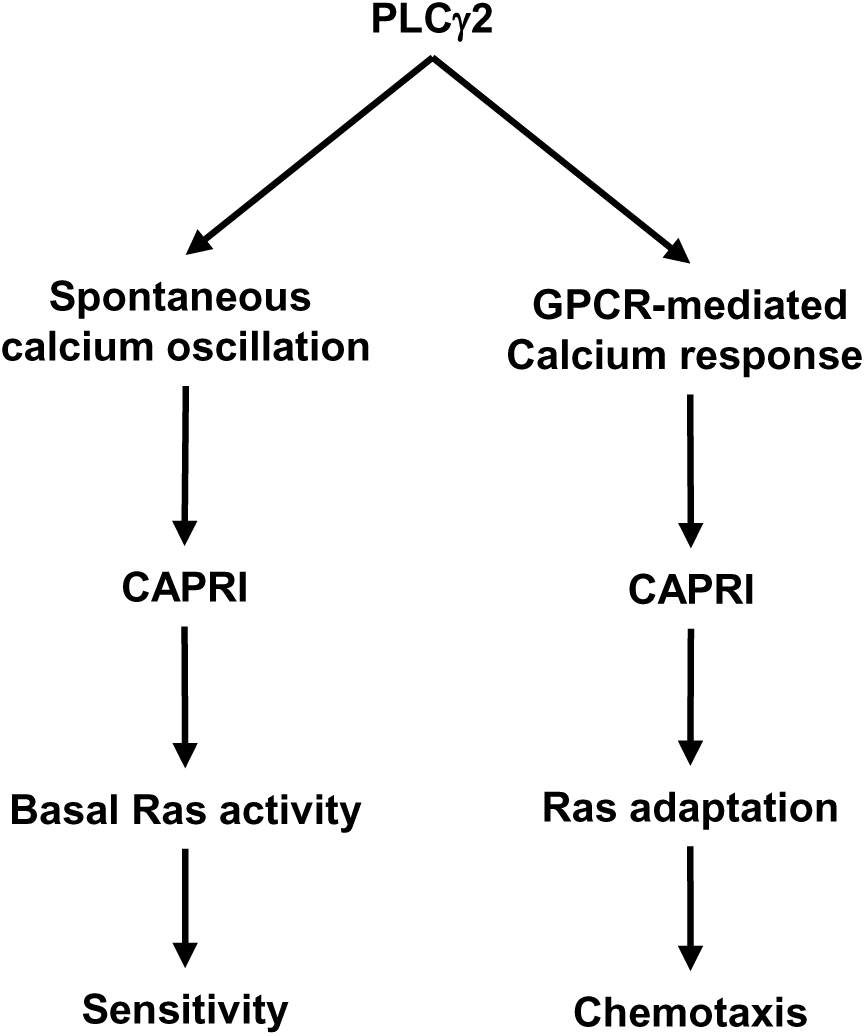
A schematic illustration of the dual roles of PLCγ2 in controlling cell sensitivity and GPCR-mediated chemotaxis.

Calcium oscillation is ubiquitous, triggered either spontaneously or upon receptor-ligand binding in all cells. The PLC-derived, IP_3_-mediated intracellular Ca^2+^ release triggers the initial [Ca^2+^] increase and constitutes calcium oscillation and calcium influx, which includes the entry of Ca^2+^ through the activation of store-operated channels (SOCs) in the plasma membrane. Murine PLCβ2/β3-deficient (*plcb2^−/−^b3^−/−^*) neutrophils display a significant decrease in IP_3_ production and calcium response, demonstrating the essential role of PLCβ2/β3 in chemoattractant-mediated calcium response (Li et al., 2000). However, mammalian neutrophils express three main isoforms of PLC, including -β2, -β3, and -γ2 (Suh et al., 2008). We previously reported that chemoattractant stimulation induces robust plasma membrane translocation of PLCγ2 (Xu et al., 2015), which is a highly expressed PLC isoform that can be activated by membrane translocation (Falasca et al., 1998; Nishida et al., 2003). We also found that the membrane translocation of PLCγ2 requires its C2-domain (Xu et al., 2023b). To delineate the contribution of PLCβ2/β3 and γ2 in chemoattractant-induced calcium signaling, we monitored calcium response in *plcg2^kd^* cells, which express endogenous PLCβ2/β3. In these cells, the chemoattractant stimulation-triggered calcium response results from the activation of PLCβ2/β3. The observed difference between CTL and *plcg2^kd^* cells is the contribution of PLCγ2 in this process. *plcg2^kd^*neutrophils display a concentration-dependent calcium response: in response to stimuli at a saturating dose, *plcg2^kd^* neutrophils display calcium responses with a normal amplitude, but with significantly decreased duration or secondary (oscillatory) calcium response (Xu et al., 2023b); upon medium (10 nM fMLP) or low (1 nM fMLP) stimuli, *plcg2^kd^* neutrophils display significantly reduced calcium response in both amplitude and duration; upon subsensitive stimuli, *plcg2^kd^* neutrophils do not display spontaneous calcium oscillation (**Figure 1**). Taken together, the above results indicate that PLCβ2/β3 is responsible for the initial calcium response upon chemoattractant stimulation, and PLCγ2 is responsible for maintaining the duration of the calcium response and mediating the spontaneous calcium oscillation.

Few connections have been made between calcium oscillation and cell sensitivity to extracellular stimuli. No clear biological function of calcium oscillation had been implicated in chemotaxis of neutrophils. In both the model organism *Dictyostelium* and mammalian neutrophils, Ras plays a central role in the signaling pathways of chemotaxis of eukaryotic cells and is the hallmark of basal cell sensitivity. Cells lacking negative regulators of Ras signaling, such as *c2gapA^−^ Dictyostelium* cells or *capri^kd^*neutrophils, often display an increased basal Ras activity and a hypersensitivity to stimuli (Xu and Jin, 2022). The consequence is an upshift in the concentration range of chemoattractant gradients, in which cells can sense and chemotax. That is, these cells are able to sense and chemotax in the gradient at subsensitive concentrations but fail to chemotax in the gradients at saturating concentrations. Plasma membrane (PM) targeting of these Ras GAPs is required for their functions and often requires calcium signaling. In neutrophils, we found that the recruitment of CAPRI to PM significant reduced in *plcg2^kd^*neutrophils in both resting and chemoattractant-stimulating states (**Figure 2**) (Xu et al., 2023b). Same as *capri^kd^* neutrophils, *plcg2^kd^*neutrophils also display an increased Ras activity, sensitivity, and an upshift in the concentration range of chemoattractant gradients (**Figure 2-7**) (Xu et al., 2023b). Our result reveals a molecular mechanism by which neutrophils employ PLCγ2 to mediate spontaneous calcium oscillation to control cell sensitivity to stimuli. In addition, neutrophils use PLCγ2 to constitute the essential calcium signaling to recruit CAPRI for Ras deactivation for a proper Ras adaptation (Xu et al., 2023b). In conclusion, neutrophils apply PLCγ2 to gate the concentration range for neutrophil chemotaxis.

## ACKNOWLEDGMENTS

This work was supported by DIR, NIAID (National Institute of Allergy and Infectious Diseases), NIH (National Institutes of Health).

## MATERIALS AND METHODS

## AUTHOR CONTRIBUTIONS

Conceptualization: X.X.; Investigation: X. X., W.K.; Data analysis: X. X., W.K., A.L.; Writing – Original draft, X.X., Review & Editing, X.X., W.K., T.J.

## SUPPLEMENTARY MATERIALS

### Supplementary videos

**Video S1-S3.** Calcium response in the control (CTL, left) and *plcg2^kd^* (right) human neutrophil-like cell HL60 upon 10 nM (**S1**), 1 nM (**S2**), or 0.1 nM (**S3**) fMLP, respectively. Cells were stained with the calcium indicator Fluo-4 and stimulated with 10, 1, or 0.1 nM fMLP at the beginning of the movies, respectively. Scale bar = 50 μm.

**Video S4-6.** Membrane translocation of CAPRI-GFP in the control (CTL, left) and *plcg2^kd^* (right) HL60 cells upon 10 nM (**S4**), 1 nM (**S5**), or 0.1 nM (**S6**) fMLP, respectively. Cells expressing CAPRI-GFP (green) were stimulated with 10, 1, or 0.1 nM fMLP at the beginning of the movies, respectively. To visualize the application of the stimuli, fMLP was mixed with a red fluorescent dye Alexa 594 (red). Scale bar = 5 μm.

**Video S7-S8.** Membrane translocation of active Ras biosensor, RBD-RFP, in the control (CTL, left) and *plcg2^kd^* (right) HL60 cells upon fMLP stimulation. Cells expressing RBD-RFP (red) were stimulated with 10 nM (**S7**) or 0.1 nM (**S8**) fMLP at the beginning of the movies. To visualize the application of the stimuli, fMLP was mixed with a green, fluorescent dye Alexa 488 (green). Scale bar = 5 μm.

**Video S9-S10.** Membrane translocation of PIP_3_ biosensor, PH-GFP, in the control (CTL, left) and *plcg2^kd^* (right) HL60 cells upon fMLP stimulation. Cells expressing PH-GFP (green) and a PM marker (red) were stimulated with 10 nM (**S9**) or 0.1 nM (**S10**) fMLP, respectively, at the beginning of the movies. Scale bar = 5 μm.

**Video S11-12**. Monitoring fMLP-induced actin polymerization in CTL and *plcg2^kd^* (right) cells upon fMLP stimulation. Actin polymerization is monitored by the membrane translocation of GFP-F-tractin in cells. 10 nM (**S11**) or 0.1 nM nM (**S12**) fMLP was added to the cells at the beginning of the movies. Scale bar = 5 μm.

**Video S13**. Montaged movie shows the movement of CTL (left) or *plcg2^kd^* (right) cells experiencing no gradient (top), or gradients generated from the chemoattractant sources at the indicated concentrations.

## REFERENCES

Behjati, S., P.S. Tarpey, H. Sheldon, I. Martincorena, P. Van Loo, G. Gundem, D.C. Wedge, M. Ramakrishna, S.L. Cooke, N. Pillay, H.K.M. Vollan, E. Papaemmanuil, H. Koss, T.D. Bunney, C. Hardy, O.R. Joseph, S. Martin, L. Mudie, A. Butler, J.W. Teague, M. Patil, G. Steers, Y. Cao, C. Gumbs, D. Ingram, A.J. Lazar, L. Little, H. Mahadeshwar, A. Protopopov, G.A. Al Sannaa, S. Seth, X. Song, J. Tang, J. Zhang, V. Ravi, K.E. Torres, B. Khatri, D. Halai, I. Roxanis, D. Baumhoer, R. Tirabosco, M.F. Amary, C. Boshoff, U. McDermott, M. Katan, M.R. Stratton, P.A. Futreal, A.M. Flanagan, A. Harris, and P.J. Campbell. 2014. Recurrent PTPRB and PLCG1 mutations in angiosarcoma. Nat Genet. 46:376–379.

Caraux, A., N. Kim, S.E. Bell, S. Zompi, T. Ranson, S. Lesjean-Pottier, M.E. Garcia-Ojeda, M. Turner, and F. Colucci. 2006. Phospholipase C-γ2 is essential for NK cell cytotoxicity and innate immunity to malignant and virally infected cells. Blood. 107:994–1002.

Clark, A.J., R. Romero, and H.R. Petty. 2010. Improved detection of nicotinamide adenine dinucleotide phosphate oscillations within human neutrophils. Cytometry A. 77:976–982.

Danielsen, S.A., L. Cekaite, T.H. Agesen, A. Sveen, A. Nesbakken, E. Thiis-Evensen, R.I. Skotheim, G.E. Lind, and R.A. Lothe. 2011. Phospholipase C isozymes are deregulated in colorectal cancer--insights gained from gene set enrichment analysis of the transcriptome. PLoS One. 6:e24419.

Falasca, M., S.K. Logan, V.P. Lehto, G. Baccante, M.A. Lemmon, and J. Schlessinger. 1998. Activation of phospholipase C gamma by PI 3-kinase-induced PH domain-mediated membrane targeting. Embo j. 17:414–422.

Jakus, Z., E. Simon, D. Frommhold, M. Sperandio, and A. Mócsai. 2009. Critical role of phospholipase Cγ2 in integrin and Fc receptor-mediated neutrophil functions and the effector phase of autoimmune arthritis. Journal of Experimental Medicine. 206:577–593.

Jing, H., A. Reed, O.A. Ulanovskaya, J.-S. Grigoleit, D.M. Herbst, C.L. Henry, H. Li, S. Barbas, J. Germain, K. Masuda, and B.F. Cravatt. 2021. Phospholipase Cγ2 regulates endocannabinoid and eicosanoid networks in innate immune cells. Proceedings of the National Academy of Sciences. 118:e2112971118.

Koss, H., T.D. Bunney, S. Behjati, and M. Katan. 2014. Dysfunction of phospholipase Cgamma in immune disorders and cancer. Trends Biochem Sci. 39:603–611.

Li, Z., H. Jiang, W. Xie, Z. Zhang, A.V. Smrcka, and D. Wu. 2000. Roles of PLC-beta2 and -beta3 and PI3Kgamma in chemoattractant-mediated signal transduction. Science. 287:1046–1049.

Liu, Q., S.A. Walker, D. Gao, J.A. Taylor, Y.F. Dai, R.S. Arkell, M.D. Bootman, H.L. Roderick, P.J. Cullen, and P.J. Lockyer. 2005. CAPRI and RASAL impose different modes of information processing on Ras due to contrasting temporal filtering of Ca2+. J Cell Biol. 170:183–190.

Lockyer, P.J., S. Kupzig, and P.J. Cullen. 2001. CAPRI regulates Ca(2+)-dependent inactivation of the Ras-MAPK pathway. Curr Biol. 11:981–986.

Martins, M., A. McCarthy, R. Baxendale, S. Guichard, L. Magno, N. Kessaris, M. El-Bahrawy, P. Yu, and M. Katan. 2014. Tumor suppressor role of phospholipase C epsilon in Ras-triggered cancers. Proc Natl Acad Sci U S A. 111:4239–4244.

Nalefski, E.A., and J.J. Falke. 1996. The C2 domain calcium-binding motif: Structural and functional diversity. Protein Science. 5:2375–2390.

Nishida, M., K. Sugimoto, Y. Hara, E. Mori, T. Morii, T. Kurosaki, and Y. Mori. 2003. Amplification of receptor signalling by Ca2+ entry-mediated translocation and activation of PLCgamma2 in B lymphocytes. Embo j. 22:4677–4688.

Obst, J., H.L. Hall-Roberts, T.B. Smith, M. Kreuzer, L. Magno, E. Di Daniel, J.B. Davis, and E. Mead. 2021. PLCγ2 regulates TREM2 signalling and integrin-mediated adhesion and migration of human iPSC-derived macrophages. Scientific Reports. 11:19842.

Ombrello, M.J., E.F. Remmers, G. Sun, A.F. Freeman, S. Datta, P. Torabi-Parizi, N. Subramanian, T.D. Bunney, R.W. Baxendale, M.S. Martins, N. Romberg, H. Komarow, I. Aksentijevich, H.S. Kim, J. Ho, G. Cruse, M.Y. Jung, A.M. Gilfillan, D.D. Metcalfe, C. Nelson, M. O’Brien, L. Wisch, K. Stone, D.C. Douek, C. Gandhi, A.A. Wanderer, H. Lee, S.F. Nelson, K.V. Shianna, E.T. Cirulli, D.B. Goldstein, E.O. Long, S. Moir, E. Meffre, S.M. Holland, D.L. Kastner, M. Katan, H.M. Hoffman, and J.D. Milner. 2012. Cold urticaria, immunodeficiency, and autoimmunity related to PLCG2 deletions. N Engl J Med. 366:330–338.

Palavicini, J.P., H.E. Miller, S. Smith, G. Campos, and S.C. Hopp. 2022. Multi-omics analyses reveal unexpected effects of PLCg2 deficiency in the brain. Alzheimer’s & Dementia. 18:e069302.

Petty, H.R. 2001. Neutrophil oscillations: temporal and spatiotemporal aspects of cell behavior. Immunol Res. 23:85–94.

Suh, P.G., J.I. Park, L. Manzoli, L. Cocco, J.C. Peak, M. Katan, K. Fukami, T. Kataoka, S. Yun, and S.H. Ryu. 2008. Multiple roles of phosphoinositide-specific phospholipase C isozymes. BMB reports. 41:415–434.

Suire, S., A.M. Condliffe, G.J. Ferguson, C.D. Ellson, H. Guillou, K. Davidson, H. Welch, J. Coadwell, M. Turner, E.R. Chilvers, P.T. Hawkins, and L. Stephens. 2006. Gbetagammas and the Ras binding domain of p110gamma are both important regulators of PI(3)Kgamma signalling in neutrophils. Nat Cell Biol. 8:1303–1309.

Suire, S., C. Lecureuil, K.E. Anderson, G. Damoulakis, I. Niewczas, K. Davidson, H. Guillou, D. Pan, C. Jonathan, T.H. Phillip, and L. Stephens. 2012. GPCR activation of Ras and PI3Kc in neutrophils depends on PLCb2/b3 and the RasGEF RasGRP4. EMBO J. 31:3118–3129.

Tang, W., Y. Zhang, W. Xu, T.K. Harden, J. Sondek, L. Sun, L. Li, and D. Wu. 2011. A PLCbeta/PI3Kgamma-GSK3 signaling pathway regulates cofilin phosphatase slingshot2 and neutrophil polarization and chemotaxis. Developmental cell. 21:1038–1050.

Tsai, F.C., and T. Meyer. 2012. Ca2+ pulses control local cycles of lamellipodia retraction and adhesion along the front of migrating cells. Curr Biol. 22:837–842.

Wen, X., T. Jin, and X. Xu. 2016. Imaging G Protein-coupled Receptor-mediated Chemotaxis and its Signaling Events in Neutrophil-like HL60 Cells. J Vis Exp.

Wessels, D., E. Voss, N. Von Bergen, R. Burns, J. Stites, and D.R. Soll. 1998. A computer-assisted system for reconstructing and interpreting the dynamic three-dimensional relationships of the outer surface, nucleus and pseudopods of crawling cells. Cell Motil Cytoskeleton. 41:225–246.

Xu, X., N. Gera, H. Li, M. Yun, L. Zhang, Y. Wang, Q.J. Wang, and T. Jin. 2015. GPCR-mediated PLCβγ/PKCβ/PKD signaling pathway regulates the cofilin phosphatase slingshot 2 in neutrophil chemotaxis. Mol Biol Cell. 26:874–886.

Xu, X., and T. Jin. 2022. Ras inhibitors gate chemoattractant concentration range for chemotaxis through controlling GPCR-mediated adaptation and cell sensitivity. Front Immunol. 13:1020117.

Xu, X., X. Wen, S. Bhimani, A. Moosa, D. Parsons, H. Ha, and T. Jin. 2023a. G protein-coupled receptor-mediated membrane targeting of PLCgamma2 is essential for neutrophil chemotaxis. J Leukoc Biol. 114:126–141.

Xu, X., X. Wen, S. Bhimani, A. Moosa, D. Parsons, H. Ha, and T. Jin. 2023b. G protein-coupled receptor-mediated membrane targeting of PLCγ2 is essential for neutrophil chemotaxis. J Leukoc Biol. 114:126–141.

Xu, X., X. Wen, A. Moosa, S. Bhimani, and T. Jin. 2021. Ras inhibitor CAPRI enables neutrophil-like cells to chemotax through a higher-concentration range of gradients. Proc Natl Acad Sci U S A. 118.

Xu, X., M. Yun, X. Wen, J. Brzostowski, W. Quan, Q.J. Wang, and T. Jin. 2016. Quantitative Monitoring Spatiotemporal Activation of Ras and PKD1 Using Confocal Fluorescent Microscopy. Methods Mol Biol. 1407:307–323.

Yi, J., X.S. Wu, T. Crites, and J.A. Hammer, 3rd. 2012. Actin retrograde flow and actomyosin II arc contraction drive receptor cluster dynamics at the immunological synapse in Jurkat T cells. Molecular biology of the cell. 23:834–852.

